# Comparative analysis of ChIP-exo peak-callers: impact of data quality, read duplication and binding subtypes

**DOI:** 10.1101/695890

**Authors:** Vasudha Sharma, Sharmistha Majumdar

## Abstract

**Background:** ChIP (Chromatin immunoprecipitation)-exo has emerged as an important and versatile improvement over conventional ChIP-seq as it reduces the level of noise, maps the transcription factor (TF) binding location in a very precise manner, upto single base-pair resolution, and enables binding mode prediction. Availability of numerous peak-callers for analyzing ChIP-exo reads has motivated the need to assess their performance and report which tool executes reasonably well for the task.

**Results:** This study has focussed on comparing peak-callers that report direct binding events with those that report indirect binding events. The effect of strandedness of reads and duplication of data on the performance of peak-callers has been investigated. The number of peaks reported by each peak-caller is compared followed by a comparison of the annotated motifs present in the reported peaks. The significance of peaks is assessed based on the presence of a motif in top peaks. Indirect binding tools have been compared on the basis of their ability to identify annotated motifs and predict mode of protein-DNA interaction.

**Conclusion:** By studying the output of the peak-callers investigated in this study, it is concluded that the tools that use self-learning algorithms, i.e. the tools that estimate all the essential parameters from the aligned reads, perform better than the algorithms which require formation of peak-pairs. The latest tools that account for indirect binding of TFs appear to be an upgrade over the available tools, as they are able to reveal valuable information about the mode of binding in addition to direct binding. Furthermore, the quality of ChIP-exo reads have important consequences on the output of data analysis.

## Introduction

Chromatin immunoprecipitation (ChIP) followed by DNA sequencing has been widely used to study DNA binding proteins (1–2). Recent advances in sequencing technologies accompanied with a decrease in the cost of sequencing, has revolutionized the field of genomics (3).

In the last decade, variations introduced to the traditional ChIP-sequencing protocol (4) have led to the development of modified methods like ChIP-exo (5), ChIP-Nexus (6), ATAC-seq (7), Mnase-seq (8–9), etc. This, in turn, has led to an upsurge in computational biology tools specially built for each modified method. omicX (10) reports a total of 107 software just for peak-calling; this illustrates the diversity of choices available for a single task in ChIP-seq analysis. Numerous studies have been done to compare ChIP-seq peak calling methods (11–15), but to the best of our knowledge, no such study is available for ChIP-exo peak-callers. This study focuses on the available tools for ChIP-exo peak-calling.

### Difference between ChIP-seq and ChIP-exo

The major difference between ChIP-seq and the ChIP-exo (5) sequencing method developed in 2011 by Rhee & Pugh *et. al.* (5) is the use of lambda exonuclease. The 5’ - > 3’ digestion activity of lambda exonuclease degrades unbound double-stranded DNA in the 5′-3′ direction until degradation is blocked at the border of the cross-linked protein-DNA complex. This leads to the generation of highly accurate protein-DNA footprints, up to a single nucleotide resolution.

The accumulation of reads on one particular genomic location, which is mostly treated as an amplification artifact in the case of ChIP-seq analysis, may actually be a signal in the case of ChIP-exo, where the 5’ border of reads represents the binding location of the protein under investigation. ChIP-exo reads are much shorter (30-100bp) when compared to ChIP-seq reads (150-250bp) and the corresponding libraries are less complex. In all the ChIP-exo reports till date, the peaks are more specific since they are called within a range of approximately 5bp of the actual binding site whereas, in ChIP-seq, identified binding sites are less specific since they are mapped to within ±300 base pairs, due to the heterogeneous nature of sheared DNA fragments. Further, due to higher resolution and reduced background, the required depth of sequencing coverage is much lower.

ChIP-exo bears several advantages over conventional ChIP-seq, which include (i) reduction of background noise due to non-specific genomic DNA (ii) precise identification of DNA binding location of TFs with high spatial resolution (iii) reveals different modes of TF binding including the oligomeric state of the protein (iv) can distinguish clustered binding events, which otherwise appear as a single peak (v) gives information about enrichment of other proteins that interact with the targeted protein (16–17) (vi) Proteins that are weakly bound to DNA can be detected using peak shape based algorithms (25). Despite all the advantages of ChIP-exo stated above, ChIP-seq remains the popular choice because of low library complexity issues faced in ChIP-exo. There is no proper threshold for ChIP-exo reads, beyond which they might be declared redundant.

Computational tools available for analyzing ChIP-seq data are deemed unsuitable for reaping the benefits of ChIP-exo because they are designed for longer reads and do not account for the extra digestion step in the ChIP-exo protocol that leads to much smaller reads (18–19). Several tools (eg. Exoprofiler (17), Genetrack (20), GEM (18), MACE (21), Peakzilla (22), CexoR (23) developed by different research groups have been used over the years to analyze the reads generated in a ChIP-exo experiment. A list of key features of all the tools used in this study is enlisted in Table 1. We have omitted CexoR from this analysis because of its limitations of analyzing samples with low sequencing read depth and coverage (23).

**Table 1:** Peak-callers used for comparison in this study along with their key features and output formats.

| Tool | Key feature | Output |
| --- | --- | --- |
| MACS (27) | 1. Uses bimodal distribution of reads to model fragment length<br>2. Uses dynamic Poisson distribution to compare test and control samples | 1. Peak position<br>2. p-value (based on pileup height at peak summit) and q value (against random Poisson distribution with local lambda) |
| GEM (18) | 1. Uses a generative probabilistic model to assign positions to the reads after each iteration<br>2. Reciprocally links binding event discovery and motif discovery<br>3. Resolves closely spaced binding events | 1. Binding events file (including location, IP strength, fold enrichment, p-value is computed from the Binomial test when control data is available, p-value computed from Poisson test in the absence of control data, divergence of the IP reads from the empirical read distribution, fraction of noise, KmerGroup and p-value associated to the K-mer and strand),<br>2. Motif files<br>3. K-mer set memory motifs<br>4. HTML output<br>5. Read distribution file<br>6. The spatial distribution between primary and secondary motifs |
| Peakzilla (22) | 1. Estimates all parameters from the data itself<br>2. Uses bimodal distribution of reads to calculate fragment length and predict binding sites<br>3. Resolves closely spaced binding events | 1. Peak file with exact position, summit, score (based on read distribution in peaks that fits bimodal tag distribution and chi-square test), FDR, fold enrichment.<br>2. Negative peaks in the presence of control. |
| Genetrack (20) | 1. Rapid data smoothing using Gaussian smoothing<br>2. Peak detection by selecting the highest peak in a local maximum with an exclusion zone of up to a few hundred bp<br>3. Combines strand information in a composite value<br>4. Requires manual pairing of border peaks | 1. Gff file with chromosome, peak exclusion zone, tag sum, strand information and standard deviation of reads in the peak exclusion zone |
|  | 1. Normalizes and corrects sequencing data for any biases<br>2. Consolidates signal to noise ratio by | 1. BED file containing border pairs of the binding event, the method for |
| MACE (21) | reducing noise<br>3. Detects border peaks using the Chebyshev Inequality and pairs them using Gale-Shapley stable matching algorithm | detecting each border pair and corresponding p-value (composite p-value of two borders in a pair) |
| Exoprofiler (17) | 1. Useful to detect different types of footprints<br>2. The peaks are scanned against the motif database to find the highest scoring peaks<br>3. High scoring peaks are then used to calculate 5' ChIP-exo coverage of reads relative to the TFBS center to find the protein-DNA crosslink boundaries | 1. Heat map of 5' ChIP-exo coverage<br>2. Footprint profile of 5' coverage of all reads<br>3. Footprint profile of the 5' coverage of reads on both strands matching the scanned motif (output of motif permutation) |
| ChExMix (25) | 1. Probabilistic mixture model for characterizing different modes of DNA-protein interactions<br>2. Expectation Maximization (EM) algorithm for estimating binding subtype probability for each binding event | 1. Event subtype file (reports total read count, signal fraction, binding coordinate, fold enrichment, event subtype, binding sequence, log[2]p-value(log likelihood score of subtype specificity for a motif hit))<br>2. Motif file<br>3. Peak-peak distance histogram<br>4. Peak-motif distance histogram |

Although numerous computational tools are available, a tangible framework, which can be utilized to analyze the vast amount of complex genomic data, is still missing. There is a variety of tools available, all based on different algorithms, which makes it difficult for the end user to make a choice.

In this study, we have compared the most popular peak callers developed for ChIP-exo reads based on the (i) number of binding events reported and (ii) motif discovery from the peak output. We have implemented the above-listed tools on publicly available ChIP-exo data of glucocorticoid receptor (GR) from three different cell lines (17) to draw an objective comparison among the available software. Since these tools are mostly used by biologists with limited computational experience, we have compared the tools using default parameters to conclude which one works best without any modifications. We have also tried to investigate how the quality of the experiment performed, affects the peak calling step of the analysis.

### Classification of ChIP-exo tools

Due to variations in the output of these tools, it is hard to assess their performance using the same parameters. So for the ease of understanding, we have classified the available tools into two broad categories:

#### 1. Tools that report binding event subtypes

These tools give a protein-DNA crosslinking pattern using ChIP-exo tag distribution, which can be further used to classify binding event subtypes of the protein of interest. ExoProfiler and ChExMix fall in this category.

ExoProfiler analyses the ChIP-exo signal for alternative modes of TF binding. It scans the peaks for motifs given that the binding motif is already known. It then finds the 5’ read coverage around the motif to discover the type of binding. ChExMix, on the other hand, utilizes a probabilistic mixture model to use sequencing tag enrichment patterns and DNA motifs for TF binding; unlike ExoProfiler, it does not require a subtype binding event to contain a motif instance. ChExMix is also capable of *de novo* motif discovery using MEME (26) and can use peaks reported by other peak callers as input.

#### 2. Tools that report direct binding events

These tools follow the classical approach of signal enrichment by an accumulation of reads over a genomic location. GEM, MACE, Genetrack, Peakzilla fall into this category.

GEM uses an empirical distribution of ChIP-exo reads to identify the cross-linking pattern and if the distribution is not specified, it automatically learns a model from sequences around binding events. MACE outputs a border pair of binding positions where right and left borders denote the 5’ position of TF binding on top and bottom strands respectively. Genetrack requires post-processing of peak files for ChIP-exo samples; here the peaks which are a fixed distance apart (distance is selected by the user depending on factors including sonication fragment length and an idea of the length of protein footprint) on opposite strands are selected to form a peak-pair similar to the border pair in MACE. Peakzilla is similar to MACS and works better for TFs with narrow peaks.

We have also incorporated MACS2 (27) in this analysis to get an idea of its performance with ChIP-exo samples.

### Why GR dataset?

GR is known to have a broad spectrum of binding sites, including canonical GBS (GR binding site) as well as binding by protein-protein interactions via recruiting other TFs eg: FOX, JUN (17, 24). The three cell lines IMR90, K562, and U2OS reportedly (17) have different GR binding profiles; in IMR90 data the binding loci are shared by GR and STAT proteins; it also reveals a high number of binding motifs for FOX proteins. K562 cells are enriched in GATA binding sequences while U2OS ChIP-exo data is highly enriched with GBS (17). In this study, an attempt is made to check if different peak callers can successfully report these motifs in GR ChIP-exo data.

## Results

### Strandedness and duplication of reads influence the peak calling of ChIP-exo data

ChIPexoQual (28) is extremely useful to determine sample quality before proceeding with the analysis. It gives the user an idea about how well the experiment has performed based on the library complexity, enrichment, and strandedness. IMR90 and K562 datasets are highly duplicated with approximately 86% and 93% redundancy rates respectively, as compared to the U2OS dataset which has a low 24% redundancy rate. Although reads are expected to accumulate over a few genomic locations in ChIP-exo samples, it is hard to distinguish whether this accumulation of reads is due to signal or PCR artifacts. The suitability of reads to be analyzed by ChIP-exo peak-callers can be estimated by FSR (Forward Strand Ratio) plots reported by ChIPexoQual, which gives a fair idea of strandedness of reads, which in turn is an important measure in many algorithms specific to ChIP-exo data (GEM, MACE, CexoR, Genetrack, ExoProfiler).

In the ChIP-exo datasets from the IMR90 and U2OS cell types, Unique Read Coefficient (URC) is very high and it decreases with increase in Average Read Coefficient (ARC), implying high ChIP enrichment and library complexity (28) (Fig. 1a). The same trend is observed in K562 cell line indicating high ChIP enrichment; however, the URC value is low in comparison to other cell types which suggests that the K562 library is less complex than the other two samples (Fig. 1a).

**Fig.1.**
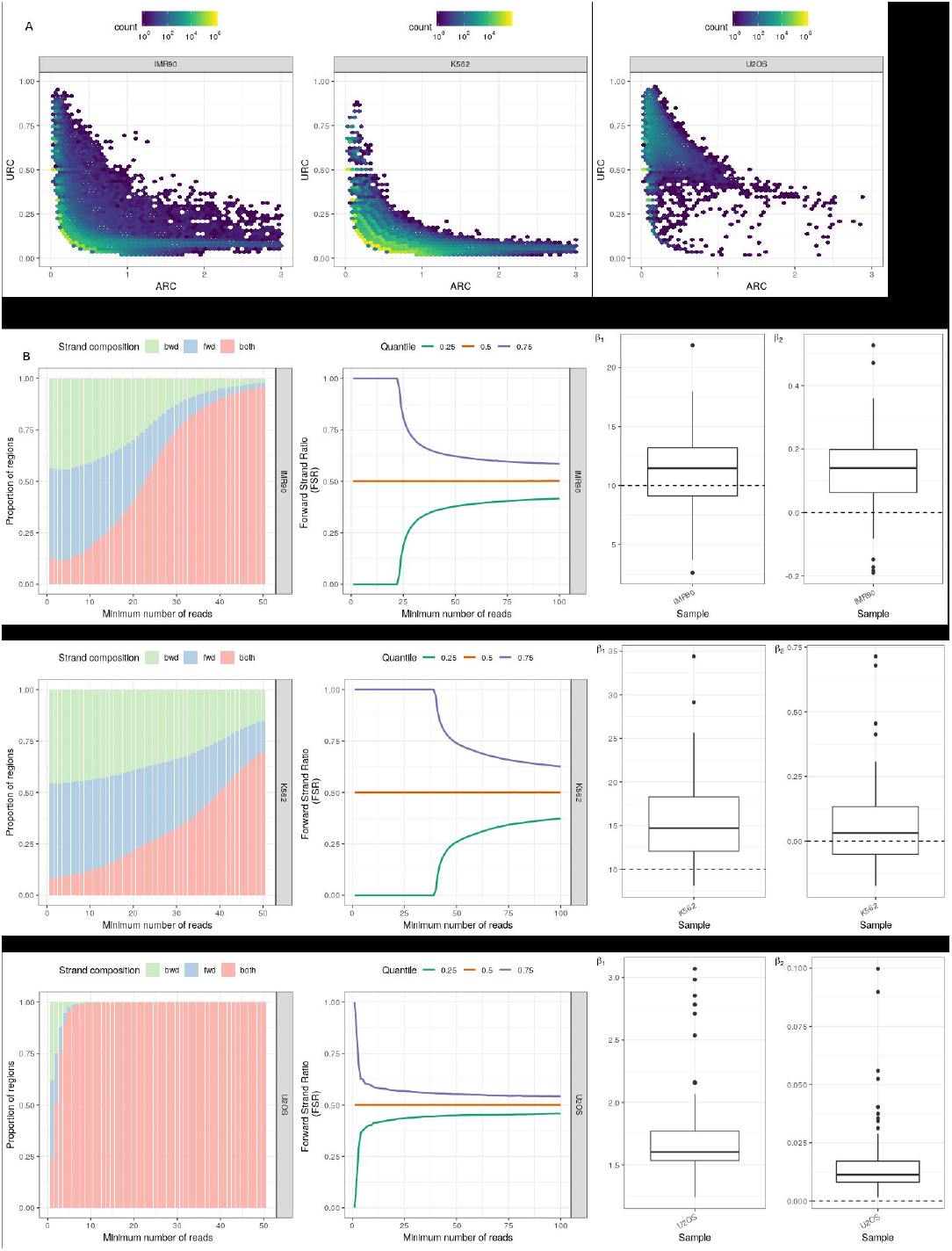
Quality metrics of ChIP-exo datasets as reported by ChIPexoQual 1a) ARC vs. URC plots for IMR90, K562 and U2OS datasets. The color represents the number of read islands (enriched regions) or bins, and with increasing number of read islands, the color shifts from blue to yellow 1b) Region composite plots and Forward Strand Ratio plots for IMR90, K562 and U2OS datasets representing the strand compositions of read islands (enriched regions). Left panel: Region composite plots, in which green represents the proportion of read islands that have reads only on the reverse strand, blue represents the proportion with reads on forward strands and red represents read islands with reads on both the strands. Right panel: FSR plots in which quantiles are marked with green (0.25), red (0.5) and purple (0.75) 1c) β1 and β2, estimates of library complexity for IMR90, K562 and U2OS datasets. The box and whiskers plot here, gives the median value of β1 and β2 for all three cell types.

Ideally performed ChIP-exo experiments show exponential decay in the proportion of singlestranded regions in Region composite plots. More than half of the reads in IMR90 and K562 datasets fall only on a single strand while in the U2OS dataset, maximum reads fall on both the strands. The U2OS sample demonstrates a decrease in the proportion of reads on a single strand whereas in IMR90 and K562 cell samples, the proportion of reads on a single strand are more spread around the median value (Fig. 1b, left panel).

In a well performed ChIP-exo experiment, the enriched regions are expected to have an equal number of reads on forward and reverse strand. FSR plots (Fig. 1b) depict the rate at which the FSR value reaches 0.5 (indicates that there is an approximately equal number of reads on both strands). The FSR value quickly reaches 0.5 in both the quantiles for the U2OS sample as compared to both IMR90 and K562 cells. The constant value of FSR for a low minimum number of reads in IMR90 and K562 datasets indicates that these samples have very few unique positions to which reads are aligned. In other words, both these samples are highly duplicated. K562 is more duplicated than IMR90. On the other hand, the FSR plot (Fig. 1b) for U2OS cells has an approximately equal number of reads on forward and reverse strands, which implies an equal distribution of reads, as desired in an ideal ChIP-exo experiment.

The final step of the ChIPexoQual pipeline includes fitting the data to a linear model and estimate β 1 and β2 which are parameters to estimate the library complexity. Samples with β1 ≤ 10 and β2 ≃ 0 are considered as deeply-sequenced high-quality samples. The median values of β1 in both IMR90 and K562 datasets are higher than 10 whereas in U2OS dataset, it is lower than 10. The median values of β2 are higher than 0 for all three datasets. High values of β1 and β2 in IMR90 and K562 datasets (Fig. 1c) imply low library complexity and poor quality of ChIP-exo samples (28) which are not deeply sequenced.

Overall, the U2OS sample has high-quality ChIP-exo data, as compared to the other two datasets, since it has low redundancy, high ChIP enrichment, and library complexity and an approximately equal number of reads on opposite strands. Due to the huge amount of strand imbalance in IMR90 and K562 cells, it becomes difficult to identify precise border pairs of the protein-DNA crosslink pattern. In such cases where the strand imbalance is high and the libraries are less complex, the ChIP-exo samples should be analyzed like ChIP-seq experiments.

### Deduplication of reads affects the performance of ChIP-exo peak callers

The Genetrack peak-caller (represented by the number of peak-pairs and not peaks) reports the highest number of peaks in IMR90 and K562 cell types (which are highly duplicated at ∼86% and ∼93% respectively) followed by GEM, MACE, MACS, and Peakzilla (Fig. 2a). On the contrary GEM and MACS report maximum peaks for U2OS samples followed by Genetrack, MACE, and Peakzilla. Also, GEM reports the highest number of binding events in U2OS dataset (Suppl. Table S1). It should be noted that Genetrack reports the maximum number of peaks for IMR90 and K562 cell types where the read duplication levels are very high, whereas in case of U2OS cell type, the maximum number of peaks are reported by GEM followed by MACS (Fig. 2a).

**Fig.2.**
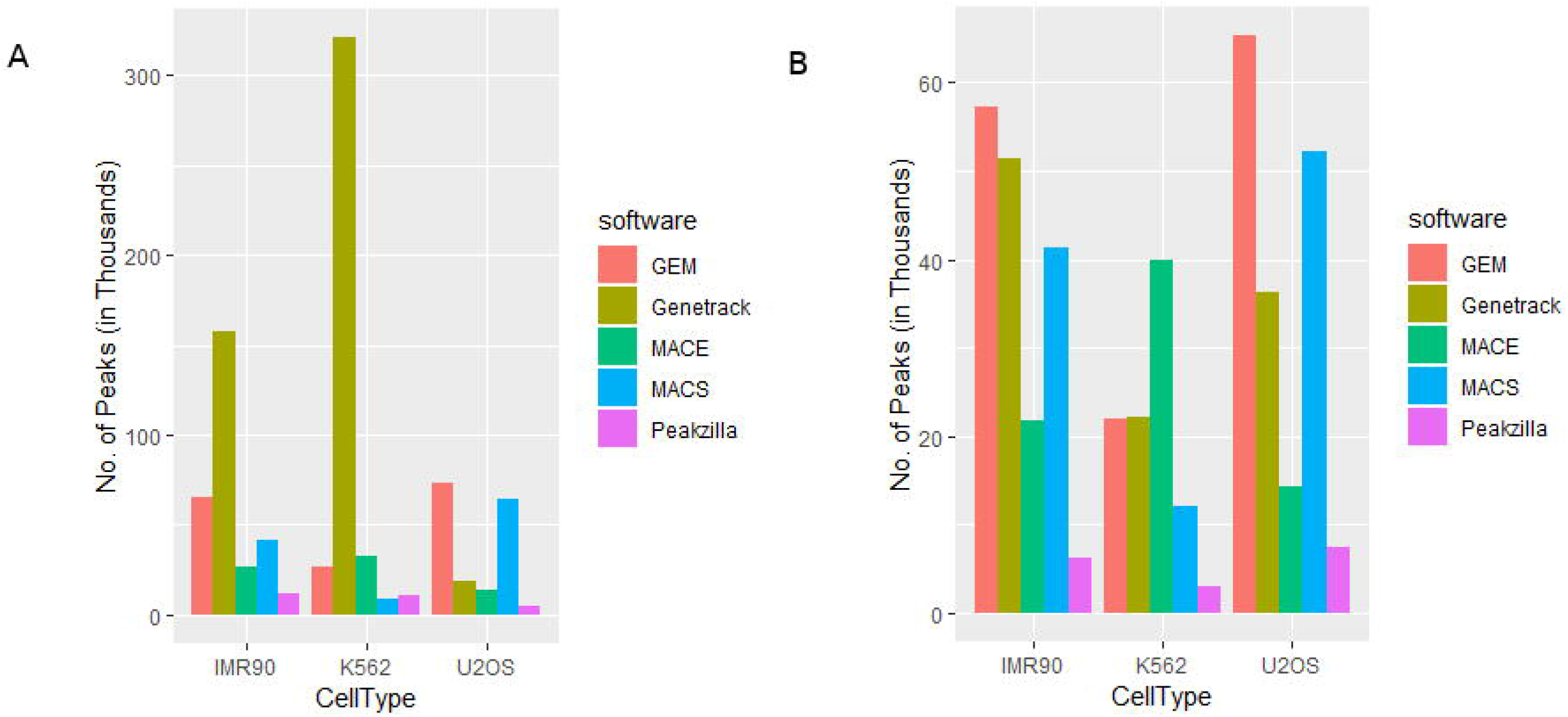
Total number of peaks/binding events reported by various peak-callers GEM, Genetrack, MACE, MACS, and Peakzilla (a) Before de-duplication of reads (b) After deduplication of reads

Genetrack identifies peaks based on a local maxima in accumulated reads; no other peak is reported within a fixed distance of the highest peak. It has no threshold of peak height beyond which a peak should be considered as a true peak (13) due to which it is capable of reporting the maximum number of peaks (Suppl. Table S1). It should also be noted that the total number of reads in the U2OS sample, as reported in the original study (17), was far less in comparison to the other cell types, due to which the number of peaks reported by each tool is less in comparison to the other two cell types.

Carroll *et al* 2014 (29) reported that the removal of duplicated reads (i.e., removal of PCR duplicates) in ChIP-exo may lead to a loss of signal. To check whether removing duplicates makes any difference to the predictions about the K562 and IMR90 datasets, the reads with identical coordinates on 5’ and 3’ ends were filtered using Picard (30) and peak calling was repeated for the deduplicated reads. (Fig. 2b). GEM, MACE, and MACS report an approximately similar number of peaks after deduplication of the datasets. However, the number of binding events discovered by Genetrack and Peakzilla drop drastically in case of IMR90 and K562 cell types (Fig. 2b), indicating that the high number of peaks reported was because of duplicate reads. But contrary to an expected decrease, the number of peaks called by Genetrack and Peakzilla increased by 2-fold and 1.5-fold respectively for U2OS cell type (Suppl.Table S2).

ChExMix has an inbuilt read filter option to remove the PCR artifacts. It forces a per-base limit on read counts to reduce the number of duplicated reads (25). ChExMix discovered 58672, 39454 and 18228 binding events for IMR90, K562 and U2OS cell types respectively which is extremely high in number in comparison to the binding events reported by ExoProfiler (4496, 313, 6236 GR binding events in IMR90, K562 and U2OS respectively) (17). It is to be noted that the binding events reported by ChExMix are a total of direct binding (where the protein is bound to a canonical sequence) and indirect binding (where the binding location is degenerate).

### Peak-pairing tools are more prone to identify false peaks

To assess the number of unique regions found by each peak caller when compared to the rest, the data reported by all the peak callers were merged into a single set, except the one whose uniqueness was to be measured. The peaks of the tool, whose uniqueness was to be found, were then intersected with the set, to report the number of unique regions discovered by each software. MACE reports maximum number of unique regions in all three datasets whereas Peakzilla does not report any unique peak (Fig. 3a).

**Fig.3.**
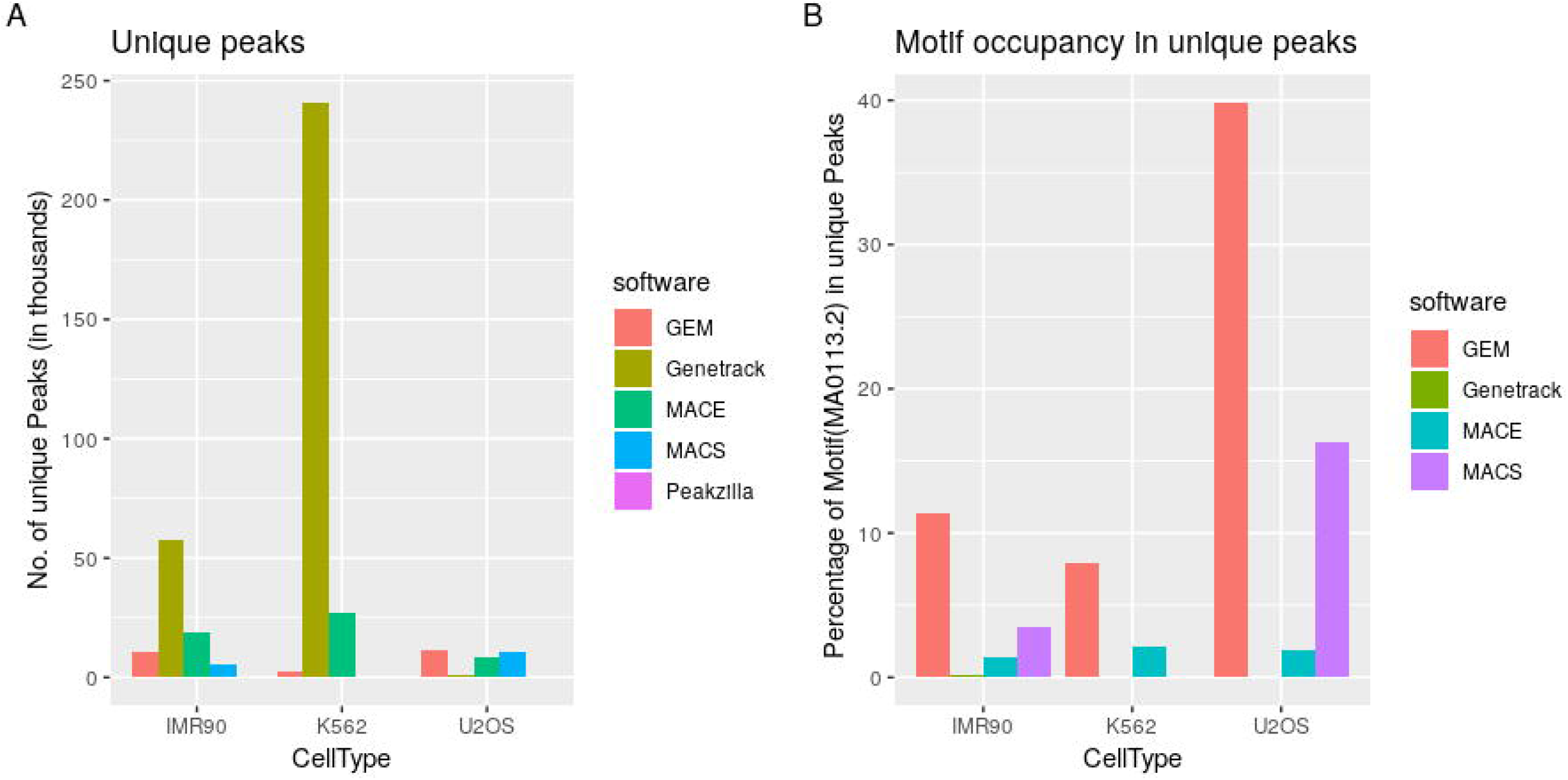
Unique regions identified by each peak caller and GBS motif occupancy in the respective unique regions. a) Number of unique peaks identified by GEM, Genetrack, MACE, MACS and Peakzilla in three cell types, IMR90, K562, and U2OS. b) Percentage of MA0113.2 motif present in the reported unique peaks as found by FIMO

These unique regions were further tested for motif occupancy (Fig. 3b). It was found that although MACE reports maximum number of unique regions, the motif occupancy in these regions is not very high. GEM, on the other hand, reported a high motif occupancy in unique peaks for all the three datasets. The maximum occupancy of GEM peaks can be explained by the fact that GEM links peak finding to motif discovery and reciprocally improves the binding event prediction around the motifs with high tag accumulation. MACE pairs the peaks on opposite strand without any constraint of a fixed sequence between the border peak-pairs. Motif occupancy in MACS peaks is also higher than MACE in IMR90 and U2OS cells but not in K562 cell type (Fig. 3b).

The unique peaks were scanned for the reported direct binding motif MA0113.2 (JASPAR (31)) for GR using FIMO (32). GEM, and MACS reported the maximum motif occupancy for the unique regions followed by MACE. Genetrack performed poorly in comparison to other tools and Peakzilla did not report any unique regions so the motif occupancy could not be found in case of Peakzilla (Fig. 3b).

The binding event predictions using tools where peak-pairs are formed by nearest peaks either by the software (MACE) or manually (Genetrack) do not perform at par with tools like MACS, Peakzilla, and GEM where most of the parameters are estimated from the data itself.

### Validation of peak caller output based on the significance score of peaks

Peaks are ranked by a significance score in the output of all peak callers and the parameters deciding the significance of a peak are different for each peak caller. The peaks with highest significance score has highest motif occupancy which decreases with the significance of peaks. To investigate which peak caller ranks the peak with highest accuracy, the peaks were sorted in descending order of their rank and top n peaks (in multiples of 50) were plotted against the fraction of annotated motifs (present within 50 bp of a peak) reported by FIMO. GEM, MACS, and Peakzilla perform consistently better in all the datasets as compared to MACE and Genetrack (Fig.4).

**Fig.4.**
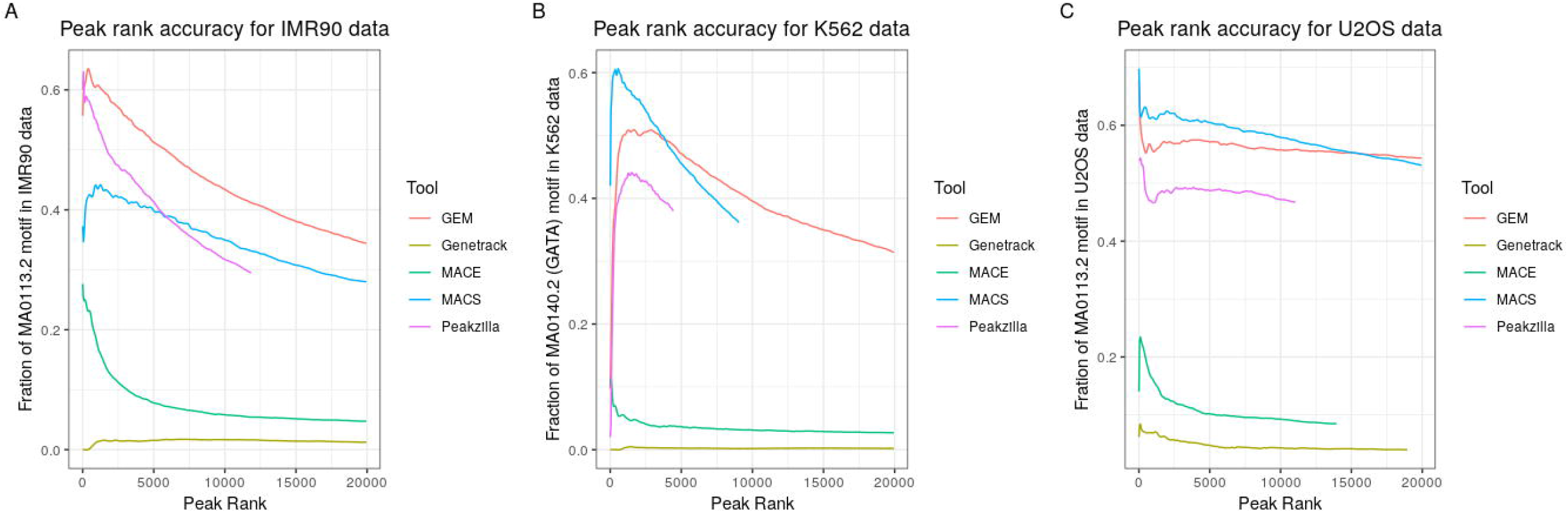
Fraction of motifs present in the top peaks of all the peak callers. a) Fraction of GBS motif in top peaks (arranged in descending order of significance score) reported in IMR90 dataset b) Fraction of GATA motif in top peaks (arranged in descending order of significance score) reported in K562 dataset c) Fraction of GBS motif in top peaks (arranged in descending order of significance score) reported in U2OS dataset.

### Total motif occupancy in peaks

GR is known to bind multiple DNA sequences via direct as well as tethered binding to DNA by binding to already recruited proteins. The peaks reported by all direct binding peak callers were scanned for the GBS motif (JASPAR MA0113.2) using FIMO (32). FIMO reported the highest number of motifs, with a p-value of less than 1e-4, in the peaks reported by the GEM output (18105 hits in IMR90, 2803 hits in K562 and 44148 hits in U2OS datasets (Table S3)) followed by MACS, Peakzilla, MACE while the least number of motifs were obtained in the Genetrack predicted binding sites (261 hits in IMR90, 158 hits in K562 and 318 in U2OS datasets (Table S3)) (Fig. 5a). The K562 dataset, which had the highest level of duplicated reads, reported the least number of motifs for all the peak callers (Fig. 5a).

**Fig.5 a.**
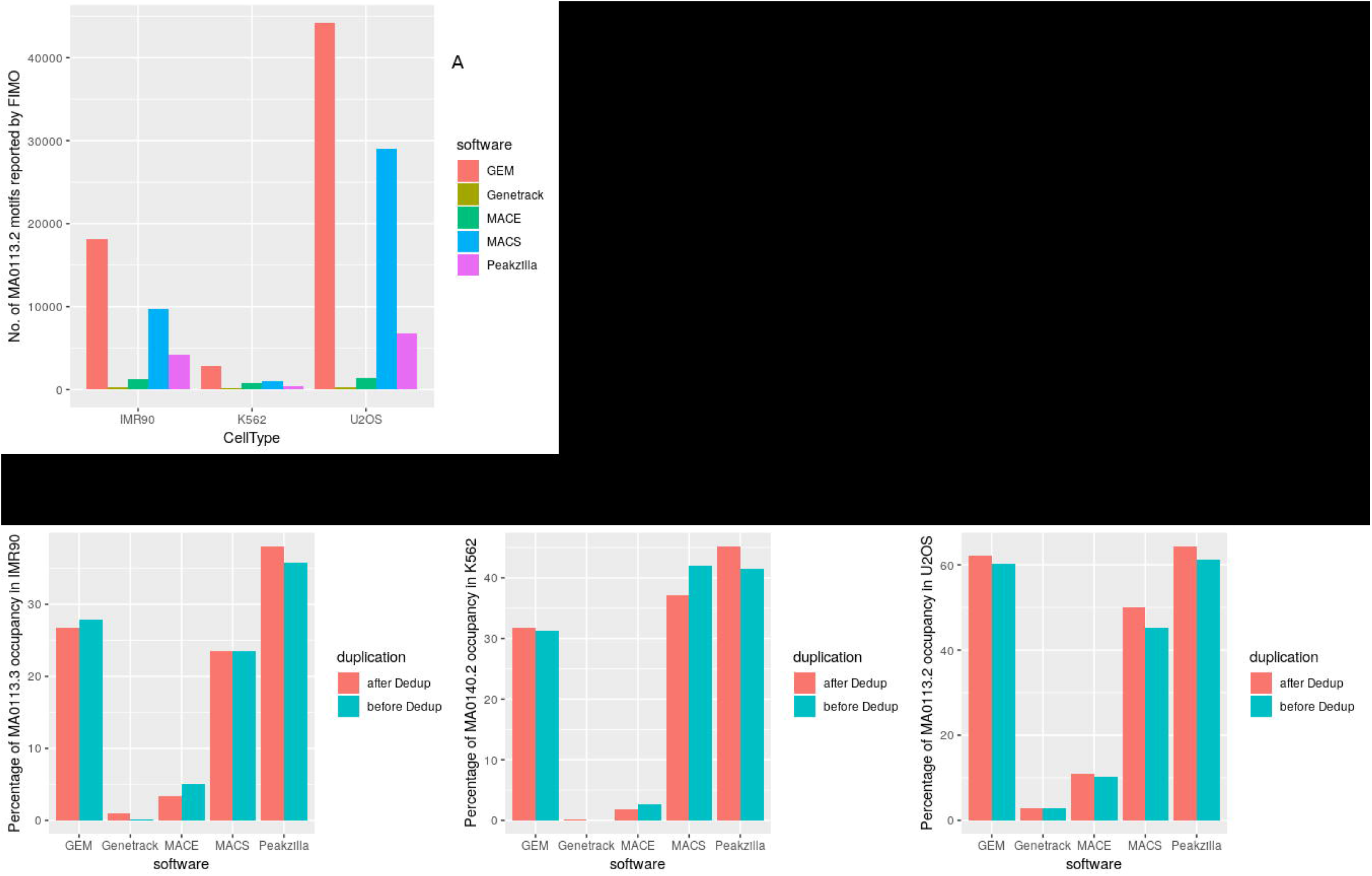
Number of GBS motifs (MA0113.2) scanned by FIMO in peak output of GEM, Genetrack, MACE, MACS, and Peakzilla.

Peaks discovered before and after deduplication were scanned for the presence of GBS, FoxA1 (JASPAR MA0148.3), GATA (JASPAR MA0140.2) and STAT3 (JASPAR MA0144.2) motifs, which have been previously reported to bind GR in multiple studies (17). When the peaks were scanned for GBS (JASPAR MA0113.2) in IMR90 and U2OS datasets and GATA (JASPAR MA0140.2) in K562 datasets (GATA sequences are known to be highly enriched in K562 cells (17)), GEM and MACS found an approximately equal number of motifs in peaks, irrespective of deduplication of reads (Fig.5b). Peakzilla identified a higher number of motifs after deduplication in the datasets thereby, implying that Peakzilla performance improves after removing PCR duplicates. A similar trend was observed when peaks were scanned for secondary motifs (FOXA1 and STAT3) in all three datasets (Suppl. Fig S1). MACE and Genetrack output had the least number of motif hits in all the datasets for all the scanned motifs including FOXA1 (JASPAR MA0148.3) and STAT3 (JASPAR MA0144.2) (Suppl. Fig S1).

**Fig.5 b.**
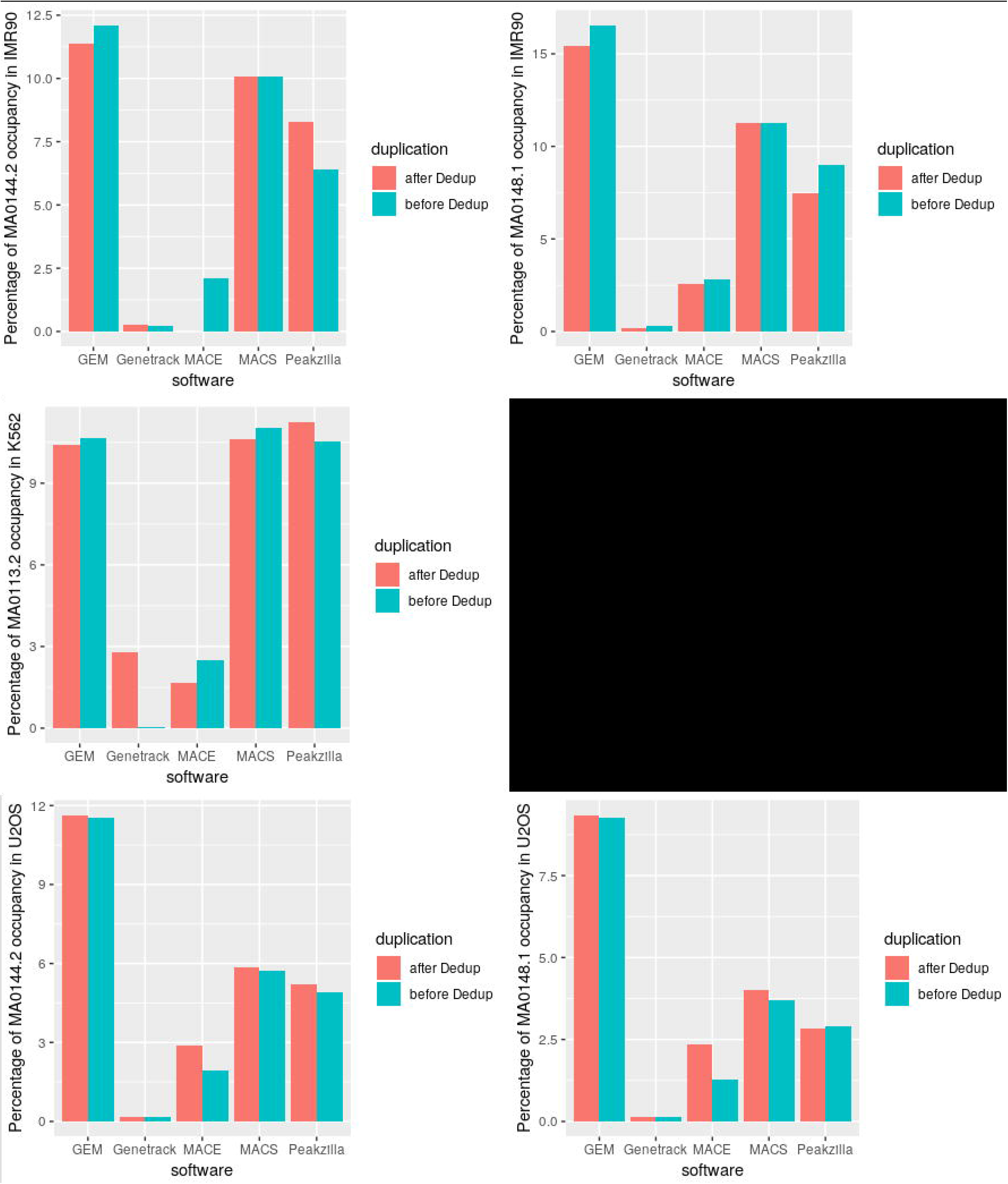
GBS (MA0113.2) and GATA (MA0140.2) occupancy in IMR90, U2OS, and K562 datasets before and after deduplication, as reported by FIMO.

Therefore, the motif occupancy reported in tools where peak-pairs are formed by nearest peaks either by the software (MACE) or manually (Genetrack) does not perform at par with tools like MACS, Peakzilla, and GEM, where most of the parameters are estimated from the data itself. GEM outperforms all the tools in the number of motifs identified in discovered peaks while minimum motif occupancy was reported in Genetrack binding events.

### *De novo* identification of TF binding site

To assess the peak-callers’ performance in terms of finding a motif that is similar to the previously reported JASPAR motif (31), the binding output from all the peak callers was submitted to MEME (26) for motif discovery. Since ChExMix and GEM has an inbuilt option for motif discovery, MEME was not used separately for motif identification for these tools.

When used with default parameters, GEM reported only half-site of GBS for both IMR90 and U2OS cell types and successfully identified GATA motif in the K562 dataset. ChExMix reported the full GBS motif, which was an exact match to the JASPAR motif, for IMR90 and U2OS datasets and the GATA sequence in the K562 data (Table 2).

**Table 2.**
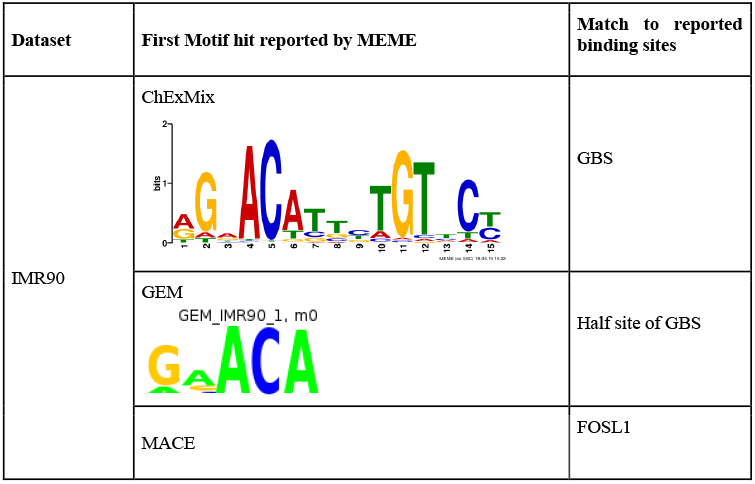
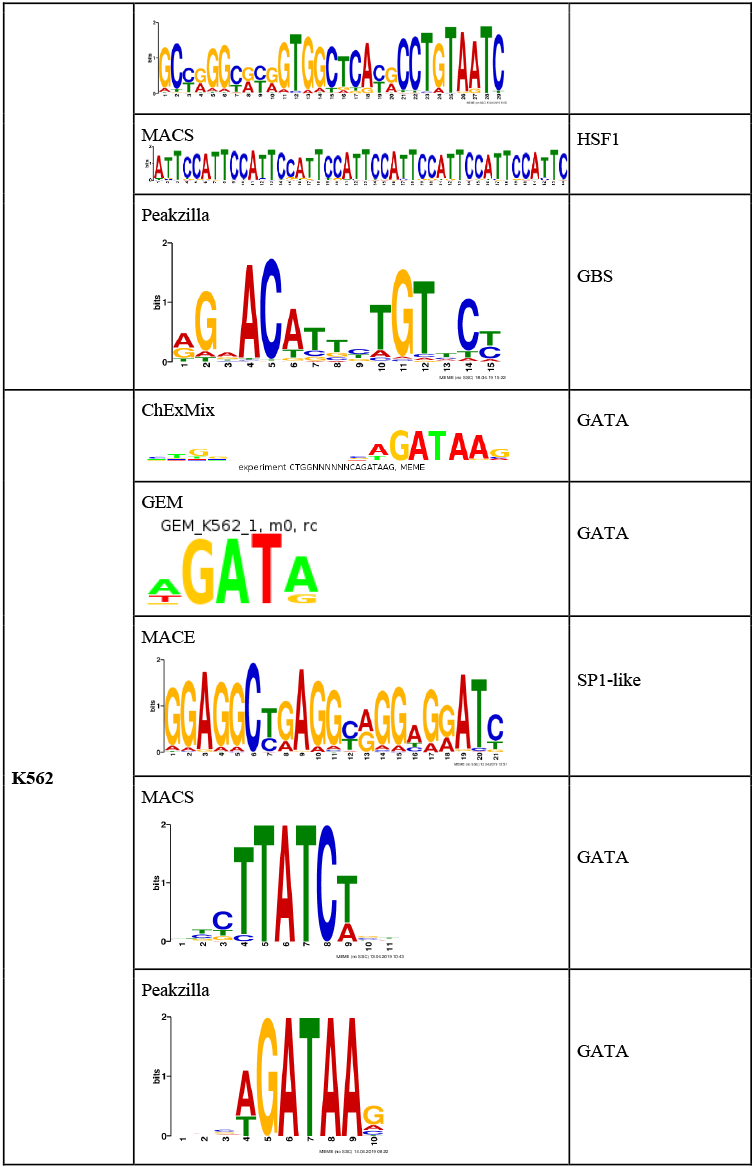
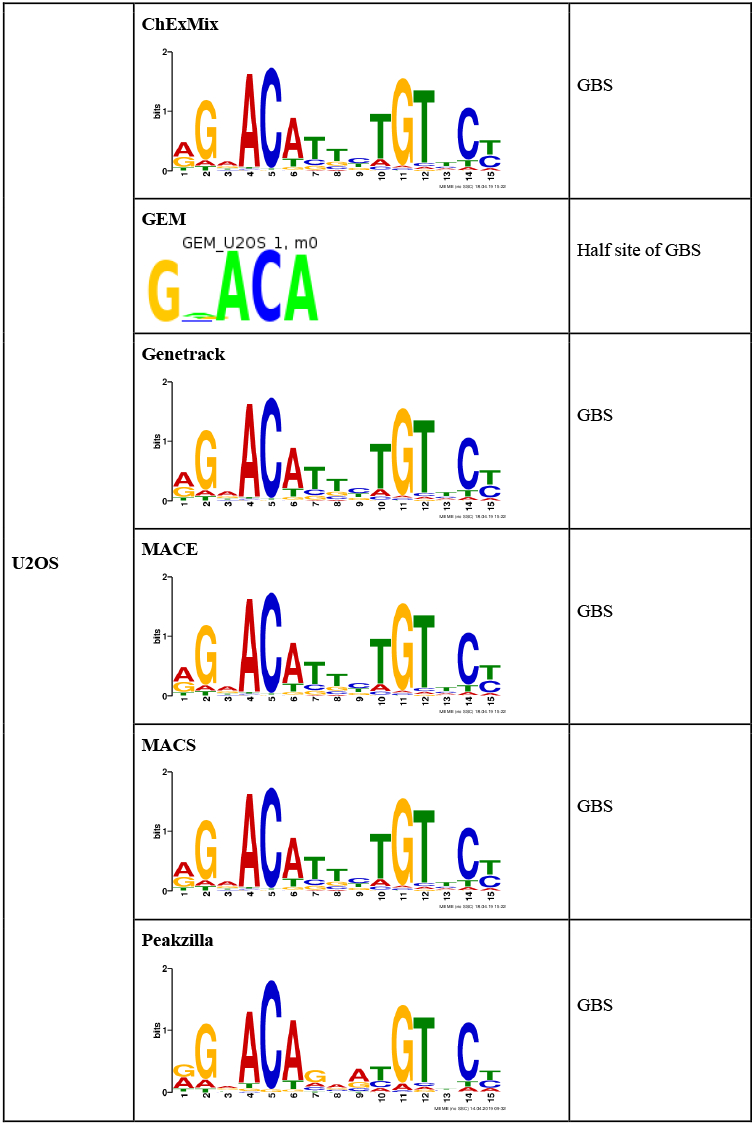
MEME output for MACE, MACS, Genetrack and Peakzilla and motifs reported directly by ChExMix and GEM.

MEME reported very long motifs for MACE peaks of IMR90 and K562 datasets, none of which matched the JASPAR motif when inspected visually and also when the motifs were submitted to TOMTOM 34) to search for similar binding sequences. Instead, FOSL1, which has been reported to be repressed by GR binding (33), was reported by TOMTOM (34) to be the best match of the motif found in MACE binding events in IMR90 dataset. In the K562 dataset, the best match for MEME motif was SP1 like sequence. GR is known to bind indirectly to SP1 sequences via SP1 TF (35). However, MEME was able to identify the GBS motif in the U2OS dataset (Table 2).

MACS identified a 44bp long repetitive motif, which resembles HSF1 binding sequence, in the IMR90 cell type. HSF proteins are known to regulate the function of GR (36). It reported GATA and GBS motifs for K562 and U2OS datasets respectively (Table 2).

MEME identified the GBS motif in the IMR90 and U2OS datasets and the GATA motif in Peakzilla peaks. In case of Genetrack, where the peaks were paired manually, MEME failed to generate a significant output for the highly duplicated IMR90 and K562 datasets but it identified the GBS motif in the U2OS data with least number of binding sites reported amongst all the tools (Table 2).

### ChExMix identifies motif with higher accuracy and mode of binding using read distribution

It is well-known that protein-DNA interactions do not depend strictly on the availability of a canonical binding motif. Proteins can interact with DNA indirectly, i.e., via protein-protein interactions or they may have a broad spectrum of recognition sequences to which they bind with a different affinity (17, 25). Starick *et al.* (17) used ChIP-seq of GR for *de-novo* motif discovery and then used these motifs along with ChIP-exo read distribution in ExoProfiler to find the binding footprint of GR and its interaction with other proteins. ChExMix encompasses both the steps in a single tool, without the requirement of a known motif.

ChExMix reported one direct binding subtype (12021 binding events with a canonical GBS site) and four indirect binding subtypes in the IMR90 dataset. Subtype 0 binds directly to GBS whereas no motif sequence is reported for the rest of the subtypes (Fig.6a). The subtype-specific read distribution profiles of subtypes 1 and 3 and subtypes 2 and 4 are very similar but there is a difference in the number of binding events (Suppl. Table S2). The tag density profiles of subtypes 1 and 3 suggest dimeric binding while that of subtypes 2 and 4 might represent composite binding. GR is known to bind a ‘combi motif’ (17). When the patterns of tag distribution profile were examined, it was found that ChExMix found the read distributions corresponding to all modes of binding reported previously for GR including, monomeric, dimeric and composite binding (Fig.6a).

**Fig.6.**
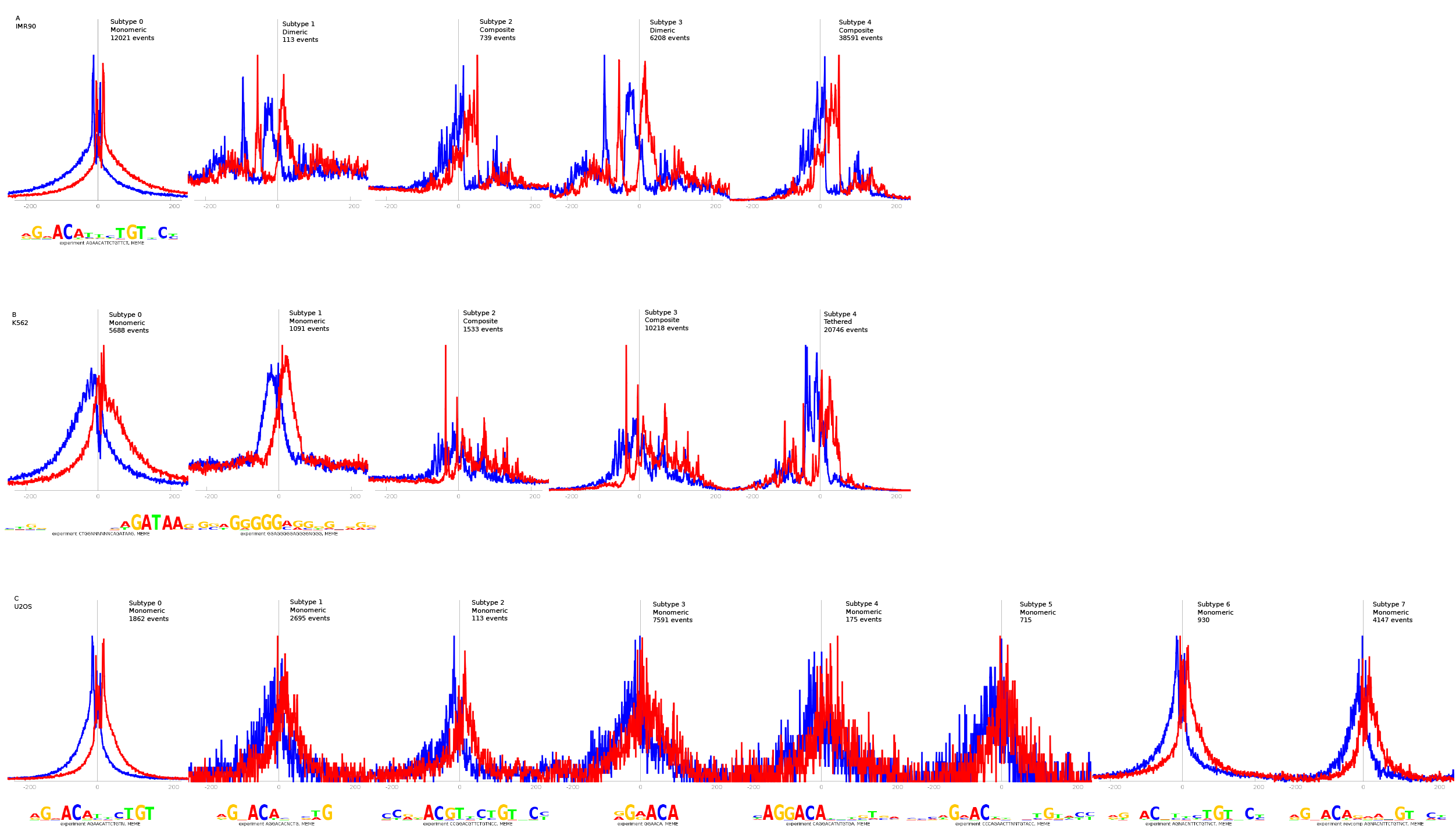
Binding subtypes discovered by ChExMix. Motifs and read distributions from (a) IMR90 dataset (b) K562 dataset (c) U2OS dataset (first subtype)

For highly duplicated datasets like K562, ChExMix filters out the duplicates based on a global per-base limit from a Poisson distribution by using a function of the number of reads and their mappability. It then estimates a permissible number of reads using the probability based on the Poisson distribution. ChExMix discovered 2 direct binding and 3 indirect binding events for the K562 dataset. Besides GATA motif, it reports SP1 binding motif (STAMP (37) E-value 6.2839e-14), a known cofactor of GR activity (38). The tag distribution profile of subtypes 2 and 3 appears to be a composite TF binding profile and that of subtype 4 appears to be tethered binding. The number of binding events of subtype 4 outnumbers all other subtypes.

ChExMix reported 7 binding subtypes, with slight differences in tag distribution profiles, for the U2OS dataset, with a total of 18228 binding events. The read distribution profile of all the subtypes is similar and each subtype has the same core GBS motif with few extra bases to the left and right-hand side of the sequence. When the intermediate output file of ChExMix was examined, it was observed that all the motifs reported by the tool were a match to the known interaction partners of GR.

## Discussion

The abundance of software available for peak-calling of ChIP-seq data has driven the need for distinguishing the available tools according to the suitability of their usage and performance. ChIP-exo is one of the modifications of ChIP-seq that has gained popularity over the years due to the precision of TF binding site detection offered by this method. This study has focused on some of the available peak-callers for ChIP-exo data, namely GEM, MACE, Genetrack, Peakzilla and recently developed ones like ExoProfiler and ChExMix. The popular ChIP-seq peak-caller MACS was also included to compare its performance on ChIP-exo data with the tools specialized for the task.

In addition to the algorithm of the peak-callers, it was observed that the quality of data highly influences the output. Clearly, the U2OS dataset, which was demonstrated to have good quality ChIP-exo data from ChIP-exoQual output, performed best with all the tools used in this study. Another important observation was that a TF can bind to different DNA sequences in different cell types.

GR is a well-characterized TF and its interaction partners are known, which made it easy to perform a comprehensible analysis of the various outputs generated from the different tools. But for *de novo* motif finding, this task would become tricky and confusing, hence there should be a proper control to differentiate true peaks from false positives. Many times, the false positives are regions that tend to be highly enriched irrespective of the ChIP experiment. For removing such biases, it is highly recommended to use an input control, if available for ChIP-exo. In the original ChIP-exo study (5), the authors were unable to generate a negative control because of the exonuclease, and for quite a long time, the use of a control was not deemed necessary for ChIP-exo experiments. But with the advancement in technologies and increasing usage of ChIP-seq, it is noted that the input control is mandatory to remove background or noise from the signal and also to reduce the discovery of false positives.

GEM appears to be the best peak finding program in direct binding tools followed by Peakzilla and MACS. Both GEM and Peakzilla can deconvolute closely spaced peaks and give a better resolution whereas MACS generates peaks with large widths, which often spans up to 200 bp (Table S4). This reduces the resolution of peaks by MACS because the resulting peaks might be a merge of multiple small peaks, which in turn makes MACS suitable for large proteins like histones. This could be the reason why MACS reported HSF1 instead of GBS motif in the IMR90 dataset. It should not be ignored that MACS, in spite of being a ChIP-seq peak calling tool, outperforms MACE and Genetrack. Peakzilla, on the other hand, identifies lesser peaks than GEM and MACS but performs reasonably well for *de novo* identification of binding motifs. It successfully reported the previously annotated motifs for all three datasets using MEME.

MACE functions by making peak-pairs of closely spaced peaks on opposite strands, while peaks need to be paired manually in Genetrack. Both MACE and Genetrack do not appear to be the best strategies to work with ChIP-exo data, because inefficient enzyme function of exonuclease and ligase affects the border formation around the protein-bound and digested DNA, which in turn influences the read accumulation around the bound protein in subsequent analysis, leading to a faulty pairing of peaks.

ChExMix and ExoProfiler use the tag distribution to identify direct as well as indirect binding. ExoProfiler has a limitation that it cannot be used for *de novo* motif discovery but is more suitable for predicting various modes of TF-DNA binding if the binding sequence is already known. ChExMix, on the other hand, overcomes this limitation by plugging into MEME to identify novel binding motifs along with the prediction of the mode of binding.

## Conclusion

In summary, this study demonstrates that the direct binding tools which learn the parameters from the reads itself (GEM, MACS, Peakzilla) serve as a better choice for peak calling ChIP-exo data over the tools which form peak-pairs to detect borders of TF binding site (MACE, Genetrack). Indirect binding tools (ChExMix and ExoProfiler) are an improvement over the existing set of tools since they also enable the user to predict the mode of binding of TF to DNA. Although the number of binding events reported by ChExMix is less than GEM, the fact that it provides an overall picture of TF binding using the shape of peak, gives it an edge over the other methods. Furthermore the quality of ChIP-exo data influences the analysis; U2OS dataset which has the best quality data out of the three cell types performs better with all the peak-calling tools.

## Methods

### Datasets

Aligned BAM files for GR ChIP-exo dataset were downloaded from EBI ArrayExpress (http://www.ebi.ac.uk/arrayexpress) under accession number E-MTAB-2956, originally submitted by Starick et. al. (17).

### Quality control

To assess the quality of the reads in GR dataset, a recently developed R package called ChIP exoQual (28) which is dedicated to quality control of ChIP-exo/ChIP-Nexus data was used. The reads were assessed for enrichment, strandedness, and complexity. ARC (average read coefficient) and URC (unique read coefficient) were used as measures to detect library enrichment and complexity and FSR (Forward Strand Ratio) plots to assess strandedness of reads.

### Peak callers

ChExMix(25), GEM(18), Genetrack(20), MACE(21), MACS(27), and Peakzilla(22) were used to call peaks using data from three cell lines. Results from ExoProfiler published in the original study (17) were used directly for comparison.

- GEM utilizes aligned read data, reference genome sequence and an empirical ChIP-exo read distribution to identify the binding events. The empirical read distribution file is used in the first step to assign priors and after the first step, the read distribution is re-estimated using the predicted binding events.
- Genetrack uses a probabilistic distribution in place of a tag and sets an exclusion zone around the mapped region to discover peak. The peaks on opposite strand, which are separated by a fixed distance, are then manually paired.
- MACE focuses on the 5’ borders of the reads, that align themselves a fixed distance apart on the reference genome, to find borders of the TF binding site. If the coverage signal at the borders of TF binding site is not high, MACE is not able to build a model for such data.
- MACS uses tag position and orientation to build a model for estimating fragment size of DNA and uses the length of fragment size to shift the tags to the 3’ end of reads. Peaks are called using the shifted tags and enrichment against background.
- Peakzilla utilizes the bimodal distribution of reads to estimate all the parameters from the aligned data to call peaks and predict the protein binding sites.
- ChExMix uses probabilistic modelling and tag distribution patterns to predict DNA-protein binding modes and binding sequence.
- ExoProfiler requires DNA binding sequence of TF along with the tag distribution patterns to identify DNA-protein binding profiles.

### Removing PCR duplicates/deduplication

Picard (30) was used to remove PCR duplicates from BAM files before another round of peak calling using GEM, Genetrack, MACE, MACS, and Peakzilla. All the tools were run using default parameters to draw a fair comparison between them. --read filter option available in ChExMix was used to remove duplicates from the datasets before running the program. Results from ExoProfiler were used directly from the original study (17).

### Unique Peaks

To identify unique peaks discovered by each peak caller, the peak output was compared against a merged dataset of the rest of the peak callers. bedtools (39) and BEDOPS (40) was used to merge the peak outputs from different peak callers and intersect it with the output of the peak caller whose uniqueness was to be measured. FIMO (32) was used to find percentage of GBS (MA0113.2) occupancy in unique peaks.

### Motif scanning

FIMO (32) was used to scan the peaks for known GBS (MA0113.2) for direct binding. Motifs with a p-value of less than 1e-4 were used to determine the occupancy in reported peaks.

### Motif discovery

The peak coordinates from all the tools except GEM and ChExMix were converted to fasta sequences using bedtools (39) and the sequences less than 8bp in size were discarded. These fasta files were then used to find enriched sequences using MEME (26). The input parameters used were as follows,

1. DNA input (fasta format)
2. Discovery mode - Classic: optimizes the E-value of the motif information content.
3. Site Distribution - zoops (one or zero motif occurrences per region).
4. Motif Count - Searching for 3 motifs.
5. Motif Width - Between 6 wide and 50 wide (inclusive).

List of tools used

1. ChIP-exoQual (version 1.8.0)
2. Picard (version 2.20)
3. ChExMix (version 0.4), GEM (version 3.4), MACE (version 1.2), MACS (version 2.1.2), Genetrack, Peakzilla
4. BEDtools (version 2.28.0), BEDOPS (2.4.36)
5. FIMO, MEME, TOMTOM,
6. STAMP
7. R (version 3.6.0)
8. GIMP (version 2.8)

## Declarations

### Ethics approval and consent to participate

Not applicable

### Consent for publication

Not applicable

### Availability of data and material

All data analyzed during this study is available in array express (See Methods section) and R script for finding peak rank accuracy using peak caller output and FIMO output has been uploaded.

### Competing interests

The authors declare that they have no competing interests

### Funding

This research was funded by IIT Gandhinagar (computers and network access required for collection, analysis and interpretation of data and in writing the manuscript; VS stipend), DBT (BT/PR16074/BID/7/569/2016) and DBT Ramalingaswami Fellowship (BT/RLF/Re-entry/43/2013) (Awarded to SM).

### Authors’ contributions

VS initialized the study and analyzed the datasets with mentioned tools. SM & VS wrote the manuscript. All authors read and approved the final manuscript.

## Acknowledgements

We thank Anirban Dasgupta for providing access to computational facilities, Gaurav Sharma for helping with custom R scripts. Jayesh Choudhari, Rachit Chhaya, and Supratim Shit for providing essential computational resources and insightful comments for this study.

## References

1. Barski A, Cuddapah S, Cui K, Roh T-Y, Schones DE, Wang Z, et al. High-resolution profiling of histone methylations in the human genome. Cell. 2007;129(4):823–37.

2. Johnson DS, Mortazavi A, Myers RM. Protein-DNA Interactions. Science. 2007;(June): 1497–503.

3. Peter J. Park. ChIP-SEQ: advantages and challenges of a maturing technology. Nat Rev Genet. 2009;10(10):669–80.

4. Furey TS. ChIP-seq and beyond: New and improved methodologies to detect and characterize protein-DNA interactions. Nat Rev Genet. 2012;13(12):840–52.

5. Rhee HS, Pugh BF. Comprehensive genome-wide protein-DNA interactions detected at singlenucleotide resolution. Cell. 2011;147(6):1408–19.

6. Qiye He, JJ. & JZ. ChIP-nexus enables improved detection of in vivo transcription factor binding footprints. Nat Biotechnol. 2015; 33:395–401.

7. Buenrostro J, Wu B, Chang H, Greenleaf W. ATAC-seq method. Curr Protoc Mol Biol. 2016;2015:1–10.

8. Zhao KC. Genome-Wide Approaches to Determining Nucleosome Occupancy in Metazoans Using MNase-Seq. Chromatin Remodel Methods Mol Biol (Methods Protoc.) 2012; 833:413–9.

9. Franklin Pugh M. Genome-Wide Mapping of Nucleosome Positions in Yeast Using High-Resolution MNase ChIP-Seq. Methods in Enzymology. 2012; 513:233–250.

10. Henry VJ, Bandrowski AE, Pepin AS, Gonzalez BJ, Desfeux A. OMICtools: an informative directory for multi-omic data analysis. Database (Oxford). 2014;2014(13): 1–5.

11. Koohy H, Down TA, Spivakov M, Hubbard T. A comparison of peak callers used for DNase-Seq data. PLoS One. 2014;9(5).

12. Steinhauser S, Kurzawa N, Eils R, Herrmann C. A comprehensive comparison of tools for differential ChIP-seq analysis. Brief Bioinform. 2016;17(October 2015):bbv110.

13. Laajala TD, Raghav S, Tuomela S, Lahesmaa R, Aittokallio T, Elo LL. A practical comparison of methods for detecting transcription factor binding sites in ChIP-seq experiments. BMC Genomics. 2009;10.

14. Szalkowski AM, Schmid CD. Rapid innovation in ChIP-seq peak-calling algorithms is outdistancing benchmarking efforts. Brief Bioinform. 2011;12(6):626–33.

15. Rye MB, Sætrom P, Drabløs F. A manually curated ChIP-seq benchmark demonstrates room for improvement in current peak-finder programs. Nucleic Acids Res. 2011;39(4).

16. Venters BJ. Insights from resolving protein-DNA interactions at near base-pair resolution. Brief Funct Genomics. 2018;17(2):80–8.

17. Starick SR, Ibn-Salem J, Jurk M, Hernandez C, Love MI, Chung H-R, et al. ChIP-exo signal associated with DNA-binding motifs provides insight into the genomic binding of the glucocorticoid receptor and cooperating transcription factors. Genome Res. 2015;25(6):825–35.

18. Guo Y, Mahony S, Gifford DK. High Resolution Genome Wide Binding Event Finding and Motif Discovery Reveals Transcription Factor Spatial Binding Constraints. PLoS Comput Biol. 2012;8(8).

19. Serandour AA, Brown GD, Cohen JD, Carroll JS. Development of an Illumina-based ChIP-exonuclease method provides insight into FoxA1-DNA binding properties. Genome Biol. 2013;14(12): 1–9.

20. Albert I, Wachi S, Jiang C, Pugh BF. GeneTrack - A genomic data processing and visualization framework. Bioinformatics. 2008;24(10):1305–6.

21. Wang L, Chen J, Wang C, Uusküla-Reimand L, Chen K, Medina-Rivera A, et al. MACE: model based analysis of ChIP-exo. Nucleic Acids Res. 2014;42(20):e156.

22. Bardet AF, Steinmann J, Bafna S, Knoblich JA, Zeitlinger J, Stark A. Identification of transcription factor binding sites from ChIP-seq data at high resolution. Bioinformatics. 2013;29(21):2705–13.

23. Madrigal P. CexoR: an R/Bioconductor package to uncover high-resolution protein-DNA interactions in ChIP-exo replicates. EMBnet.journal. 2015;21(0): 1–5.

24. Oakley R, Cidlowski J. Defence mechanisms in health and disease. J Allergy Clin Immunol. 2013;132(5): 1033–44.

25. Yamada N, Lai WKM, Farrell N, Pugh BF, Mahony S. Characterizing protein-DNA binding event subtypes in ChIP-exo data. Bioinformatics. 2019;35(6):903–13.

26. Bailey TL, Elkan C. Fitting a mixture model by expectation maximization to discover motifs in biopolymers. Proceedings Int Conf Intell Syst Mol Biol. 1994; 2: 28–36.

27. Zhang Y, Liu T, Meyer CA, Eeckhoute J, Johnson DS, Bernstein BE, et al. Model-based analysis of ChIP-Seq (MACS). Genome Biol. 2008;9(9):R137.

28. Welch R, Chung D, Grass J, Landick R, Keleş S. Data exploration, quality control and statistical analysis of ChIP-exo/nexus experiments. Nucleic Acids Res. 2017;45(15):1–14.

29. Carroll TS, Liang Z, Salama R, Stark R, de Santiago I. Impact of artifact removal on ChIP quality metrics in ChIP-seq and ChIP-exo data. Front Genet. 2014;5(APR): 1–11.

30. Picard [Internet]. Available from: http://broadinstitute.github.io/picard/

31. Mathelier A, Zhao X, Zhang AW, Parcy F, Worsley-Hunt R, Arenillas DJ, et al. JASPAR 2014: An extensively expanded and updated open-access database of transcription factor binding profiles. Nucleic Acids Res. 2014;42(D1): 142–7.

32. Grant CE, Bailey TL, Noble WS. FIMO: Scanning for occurrences of a given motif. Bioinformatics. 2011;27(7): 1017–8.

33. Lucibello FC, Slater EP, Jooss KU, Beato M, Müller R. Mutual transrepression of Fos and the glucocorticoid receptor: involvement of a functional domain in Fos which is absent in FosB. EMBO J. 2018;9(9):2827–34.

34. Gupta S, Stamatoyannopoulos JA, Bailey TL, Noble WS. Quantifying similarity between motifs. Genome Biol. 2007;8(2).

35. Ou XM, Chen K, Shih JC. Glucocorticoid and androgen activation of monoamine oxidase a is regulated differently by R1 and Sp1. J Biol Chem. 2006;281(30):21512–25.

36. Pratts WB. William B. PrattS. 1993;21455–8.

37. Mahony S, Benos P V. STAMP: A web tool for exploring DNA-binding motif similarities. Nucleic Acids Res. 2007;35(SUPPL.2):253–8.

38. Strähle U, Schmid W, Schütz G. Synergistic action of the glucocorticoid receptor with transcription factors. EMBO J. 2018;7(11):3389–95.

39. Quinlan AR, Hall IM. BEDTools: A flexible suite of utilities for comparing genomic features. Bioinformatics. 2010;26(6):841–2.

40. Neph S, Kuehn MS, Reynolds AP, Haugen E, Thurman RE, Johnson AK, et al. BEDOPS: High-performance genomic feature operations. Bioinformatics. 2012;28(14):1919–20.

