## Supplemental tables and legends for "Comparative analysis of ChIP-exo peak-callers: impact of data quality, read duplication and binding subtypes"

**Additional file**

**Table S1: Total number of peaks/ binding events discovered by each peak caller for each cell type**

|  | **IMR90** | **K562** | **U2OS** |
| --- | --- | --- | --- |
| GEM | 64900 | 26299 | 73233 |
| Genetrack (peak-pairs) | 157673 | 320945 | 19017 |
| MACE | 26117 | 32988 | 14020 |
| MACS | 41326 | 9130 | 64193 |
| Peakzilla | 11923 | 11126 | 4509 |

**Table S2: Total number of binding events discovered by each peak caller for each cell type, after filtering out the PCR duplicates**

|  | **IMR90** | **K562** | **U2OS** |
| --- | --- | --- | --- |
| GEM | 57304 | 21960 | 65277 |
| Genetrack (peak-pairs) | 51338 | 22153 | 36334 |
| MACE | 21769 | 39841 | 14247 |
| MACS | 41306 | 12010 | 52187 |
| Peakzilla | 6242 | 2951 | 7516 |

**Table S3: GBS motif hits (p-value < 1e-4) reported by FIMO**

|  | **IMR90** | **K562** | **U2OS** |
| --- | --- | --- | --- |
| GEM | 18105 | 2803 | 44148 |
| Genetrack (peak-pairs) | 261 | 158 | 318 |
| MACE | 1317 | 824 | 1423 |
| MACS | 9720 | 1009 | 28994 |
| Peakzilla | 4267 | 476 | 6800 |

**Table S4: Peak length statistics for GEM, Genetrack, MACE, MACS, and Peakzilla when run on GR ChIP-exo datasets for IMR90, K562 and U2OS cell types**

| **Tool** | **Peak length** | **Mean** | **Median** | **Minimum** |
| --- | --- | --- | --- | --- |
| GEM_IMR90 | 200 | - | - | - |
| GEM_K562 | 200 | - | - | - |
| GEM_U2OS | 200 | - | - | - |
| Genetrack_IMR90 (peak-pair distance) | 20 | - | - | - |
| Genetrack_K562 (peak-pair distance) | 20 | - | - | - |
| Genetrack_U2OS (peak-pair distance) | 20 | - | - | - |
| MACE_IMR90 | - | 49.4 | 50 | 1 |
| MACE_K562 | - | 75.8 | 78 | 1 |
| MACE_U2OS | - | 39.7 | 38 | 1 |
| MACS_IMR90 | - | 117.69 | 78 | 44 |
| MACS_K562 | - | 159.16 | 132 | 72 |
| MACS_U2OS | - | 93.95 | 72 | 41 |
| Peakzilla_IMR90 | 84 | - | - | - |
| Peakzilla_K562 | 110 | - | - | - |
| Peakzilla_U2OS | 80 | - | - | - |

Supplementary section legends:

Suppl. Fig. S1: FOXA1 (JASPAR MA0148.3) and STAT3 (JASPAR MA0144.2) occupancy in IMR90, U2OS, and GBS (JASPAR MA0113.2) in K562 datasets before and after deduplication, as reported by FIMO.

Suppl. Table S1: Total number of peaks/binding events reported by the tools in IMR90, K562 and U2OS datasets

Suppl. Table S2: Total number of binding events discovered by each peak caller for IMR90, K562, U2OS, after filtering out the PCR duplicates.

Suppl. Table S3 Total GBS motif occupancy (GBS motif hits (p-value < 1e-4) reported by FIMO)

Suppl. Table S4 Peak length statistics for GEM, Genetrack, MACE, MACS, and Peakzilla when run on GR ChIP-exo datasets for IMR90, K562 and U2OS cell types
